# Unified imputation of missing data modalities and features in multi-omic data via shared representation learning

**DOI:** 10.64898/2026.02.04.703630

**Authors:** Ananthan Nambiar, Carlo Melendez, William Stafford Noble

**Affiliations:** Department of Genome Sciences, University of Washington, Seattle, WA 98195, U.S.A.; eScience Institute, University of Washington, Seattle, WA 98195, U.S.A; Paul G. Allen School of Computer Science & Engineering, University of Washington, Seattle, WA 98195, USA

## Abstract

Multi-omic studies promise a more comprehensive view of biological systems by jointly measuring multiple molecular layers. In practice, however, such datasets are rarely complete: entire molecular modalities may be missing for many samples, and observed modalities often contain substantial feature-level missingness. Existing imputation approaches typically address only one of these two problems, relying either on feature-level imputation within a single modality or on pairwise translation models that cannot accommodate arbitrary combinations of missing modalities.

We present MIMIR, a deep learning framework for unified multi-omic imputation of bulk data that addresses both missing modalities and missing values through shared representation learning. MIMIR first learns modality-specific representations using masked autoencoders and then projects these representations into a common latent space, enabling reconstruction from any subset of observed modalities. Evaluated on pan-cancer multi-omic data from The Cancer Genome Atlas, MIMIR consistently outperforms baseline methods across a range of missing-modality and missing-value scenarios, including missing completely at random and missing not at random settings. Analysis of the learned shared space reveals structured cross-modal dependencies that explain modality-specific differences in imputation accuracy, with transcriptional and epigenetic modalities forming a strongly aligned core and copy number variation contributing more distinct signal. Together, these results demonstrate that shared representation learning provides an effective and flexible foundation for multi-omic imputation under heterogeneous patterns of missingness.

## Introduction

Diseases arise from complex interactions among molecules, pathways, and cells across biological scales. For example, genetic variants can alter transcriptional regulation and protein function, which in turn affect metabolism, interact with environmental exposures, and influence disease phenotypes. Unfortunately, standard predictive models for disease commonly rely on a single data modality, such as genomics or transcriptomics, thereby missing critical interactions between biological layers [1, 2, 3]. Even when multi-omic data are incorporated, most studies focus on pairs of modalities, such as linking gene expression to proteomics or methylation to transcriptomics, and therefore fail to capture the broader network of molecular dependencies [4, 5].

Despite increased adoption of multi-omic profiling, datasets with more than two molecular layers are rarely complete. Two distinct forms of incompleteness arise in these datasets: missing modalities, where an entire layer is absent for a sample, and missing values, where measurements within a modality are only partially observed (Figure 1a). Practical constraints such as cost, technical failures, and evolving study protocols often result in missing modalities, with many individuals assayed for only a subset of data types. Even when a modality is observed, substantial feature-level missingness is common and can vary across samples or batches. For example, DNA methylation data processing can remove unreliable loci in a sample-specific manner, resulting in substantial within-modality missingness, and some types of mass spectrometry measurements suffer from stochastic patterns of missingness. These inconsistencies make direct integration and modeling the full biological complexity underlying disease mechanisms challenging.

**Figure 1:**
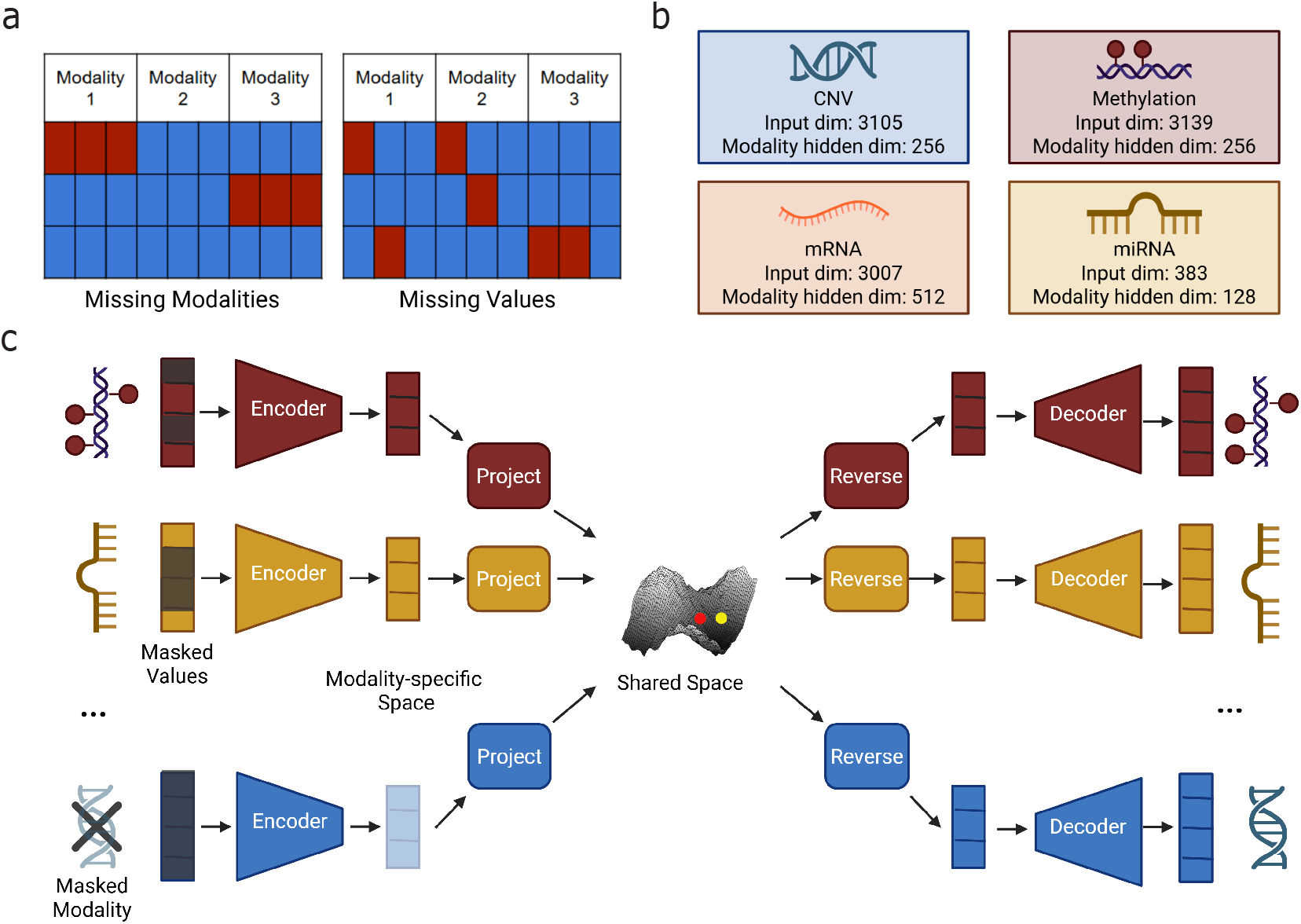
Overview of MIMIR for unified multi-omic imputation. **(a)** The two types of incompleteness commonly observed in multi-omic datasets: missing modalities, where entire modalities are absent for some samples, and missing values, where individual features within an observed modality are unmeasured. **(b)** Modality-specific autoencoders used in MIMIR, showing the input dimensionality and latent representation size for each modality. **(c)** Shared representation learning in MIMIR. Observed modalities are encoded into modality-specific latent spaces and projected into a common shared latent space. A sample representation is constructed by aggregating projected embeddings from available modalities and can be mapped back through reverse projections and decoders to reconstruct both missing values and missing modalities.

Although a recently described class of sequence-based deep learning models can predict multiple downstream molecular readouts directly from DNA sequence,[6, 7] these sequence-to-function models address a fundamentally different problem setting. They rely on a fixed genomic input and learn population-level mappings to molecular profiles, rather than modeling sample-specific variation across partially observed multi-omic datasets. As a result, sequence-to-function models are not designed to perform imputation conditioned on the subset of molecular measurements available for a given individual.

Many existing imputation approaches focus on developing feature-level imputation methods that operate within a single data type [8, 9, 10]. Although these techniques could in principle be applied to impute an entire missing data modality for a given sample, the methods are rarely designed with modality-level gaps in mind. For missing modalities, current work largely centers on pairwise translation models, which learn mappings between pairs of modalities [11, 12, 13]. More recently, several multimodal translation and integration models have been developed in single-cell settings, leveraging partially observed modalities to translate across data types [14, 15]. However, these approaches are typically tailored to specific experimental designs with limited modality combinations, and are not designed to flexibly handle arbitrary patterns of missing modalities and feature-level missingness in bulk or cohort-scale datasets. Consequently, there is, to our knowledge, no well studied unified framework for reconstructing both missing layers and missing features within those layers in large-scale bulk multi-omic settings.

To address this gap, we develop MIMIR (Multi-omic Imputation through Modality Integration and Representation learning), a cross-modal machine learning framework that integrates bulk multi-omic data and imputes missing information both across modalities and within individual modalities, enabling more complete and biologically meaningful models (Figure 1). The framework is trained in two phases. First, it is trained to learn modality-specific representations (Figure 1b). Then, it aligns the representations into a shared latent space that captures relationships spanning multiple molecular layers (Figure 1c). By jointly optimizing reconstruction, cross-modal imputation, and feature-level imputation objectives, the model can reconstruct missing layers as well as recover missing values within observed layers, accomodating the diverse and irregular missingness patterns found in real datasets (see Methods for details). We demonstrate that MIMIR accurately improves data completeness using multi-omic data from The Cancer Genome Atlas (TCGA). In doing so, this work highlights the potential for unified cross-modal imputation to enable integrative analysis and deepen our understanding of the molecular processes underlying human disease.

## Results

We evaluated MIMIR on multi-omic data from TCGA under controlled patterns of missingness. Performance was assessed by comparing imputed values to held-out ground truth across molecular modalities. We first evaluate MIMIR’s ability to reconstruct entirely missing modalities from partially observed samples and compare its performance to existing multi-omic imputation approaches under this setting. We then assess missing value imputation within observed modalities under different missingness mechanisms, including missing completely at random (MCAR) and missing not at random (MNAR), again benchmarking against baseline methods. Finally, we analyze properties of the learned shared representation to better understand how information is integrated across molecular layers.

### Missing modality imputation

MIMIR performs missing modality imputation by encoding observed modalities into a shared latent space and reconstructing the missing modality from the aggregated representation (see Methods). Performance was assessed by comparing imputed values to held-out ground truth across molecular modalities. We first evaluate MIMIR’s ability to reconstruct entirely missing modalities from partially observed samples and compare its performance to existing multi-omic imputation approaches designed for this setting. In many multi-omic studies, entire molecular modalities are missing for a subset of samples due to cost, technical failure, or study design. Recovering these missing modalities is challenging because existing approaches typically rely on pairwise translation models, which must be trained separately for each source–target modality pair and cannot naturally accommodate arbitrary combinations of observed modalities [11, 12].

MIMIR addresses this by performing cross-modal inference through a shared latent representation. At inference time, observed modalities are encoded and projected into a shared space, and their embeddings are aggregated using the same procedure as during training (Methods) to form a sample-level representation. Missing modalities are then reconstructed by decoding this shared representation through the corresponding modality-specific decoder. This inference procedure applies uniformly across all patterns of modality-level missingness and does not require retraining or modality-specific translation models.

We evaluated MIMIR’s ability to reconstruct entirely missing modalities by masking one modality at a time during evaluation and reconstructing it from the remaining observed modalities. Figure 2a shows scatter plots comparing observed and imputed values for each target modality, where each point corresponds to a held-out feature-sample pair in the test set. Imputation accuracy was quantified using Pearson correlation, computed by flattening the feature-by-sample matrix for the target modality across all test samples. MIMIR achieved strong correlations across modalities, with correlations of approximately 0.69 for CNV, 0.76 for DNA methylation, 0.78 for miRNA, and 0.78 for mRNA, indicating accurate reconstruction across diverse molecular assays. Imputed values closely track the observed values across a wide dynamic range, suggesting that the shared latent representation preserves quantitative cross-modal relationships rather than collapsing predictions toward the mean.

**Figure 2:**
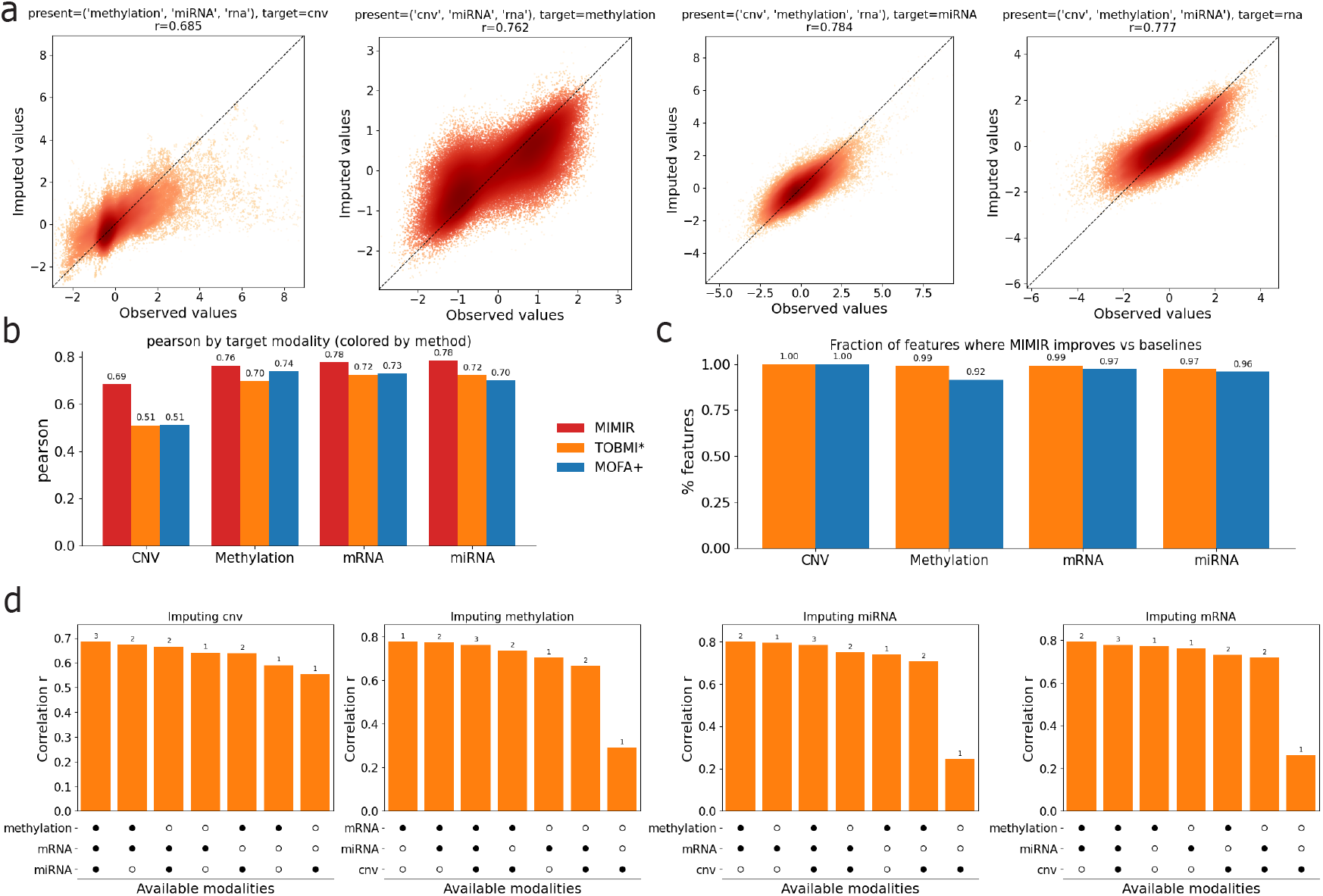
Cross-modal imputation performance of MIMIR. **(a)** Scatter plots comparing observed versus imputed values across target modalities. **(b)** Comparison of imputation accuracy (Pearson correlation) across baseline methods for each target modality. **(c)** Fraction of features for which MIMIR achieves higher correlation than baseline methods. **(d)** Imputation performance (Pearson correlation) for each target modality across different combinations of missingness. Bars are sorted by performance, and the accompanying dot matrices indicate which modalities are observed in each scenario. Numbers above bars denote the number of available modalities.

We next benchmarked MIMIR’s performance against baseline multi-omic imputation methods, including MOFA+, a latent factor model that learns a shared low-dimensional representation across modalities, and TOBMI*, a *k*-nearest-neighbor cross-modal imputation approach that reconstructs missing modalities using sample similarity in available modalities [16, 17]. Because MOFA+ does not provide a built-in procedure for missing modality prediction, missing modalities were reconstructed via post hoc projection into the learned latent space followed by reconstruction using the learned loading matrices (Methods). Figure 2b summarizes imputation accuracy for each target modality across methods. MOFA+ outperformed TOBMI* for CNV, DNA methylation, and mRNA, whereas TOBMI* achieved higher accuracy for miRNA. Across all target modalities, MIMIR achieved the highest correlation, demonstrating improved reconstruction accuracy relative to both baseline approaches. Notably, the performance gain of MIMIR was largest for CNV, where it substantially outperformed both MOFA+ and TOBMI*, indicating that the shared representation is particularly effective for reconstructing copy number variation profiles.

To assess whether these improvements reflected broad gains rather than performance driven by a small subset of features, we also examined imputation accuracy at the feature level. Figure 2c shows the fraction of features for which MIMIR achieved higher correlation than each baseline method. Across all modalities, MIMIR outperformed both MOFA+ and TOBMI* for the majority of features, indicating that its improved performance is widespread rather than confined to a narrow subset of molecular measurements.

We next examined the relationship between training sample size per subtype and reconstruction accuracy (Supplementary Figure 6). We find no significant association between subtype sample size and performance across modalities, suggesting that improvements are not primarily driven by subtype-level signal. These results indicate that MIMIR captures cross-modal relationships that extend beyond coarse clustering structure.

Finally, we examined how missing modality reconstruction depends on the types of observed input modalities. Figure 2d summarizes imputation performance for each target modality across different subsets of available modalities (indicated below each bar). Rather than improving monotonically with the number of observed modalities, reconstruction accuracy depends strongly on which modalities are present. For imputing methylation and miRNA, having mRNA available is most critical, and adding CNV provides little benefit and is slightly detrimental in the best-performing combinations. For imputing mRNA, both methylation and miRNA contribute substantially, with the highest performance observed when both are present. For imputing CNV, mRNA is again the most informative modality, with methylation and miRNA providing smaller additional gains. Together, these results demonstrate that MIMIR flexibly adapts to heterogeneous missingness patterns and that reconstruction accuracy is driven by specific cross-modal dependencies between molecular assays.

### Missing value imputation

In addition to missing entire modalities, multi-omic datasets frequently contain missing values within otherwise observed modalities. We therefore evaluated MIMIR’s ability to impute missing values within a particular modality. For missing value imputation, individual feature values were masked within observed modalities at evaluation time, while all modalities remained present. Unlike missing modality imputation where aggregation was done using a simple average, shared representations in the missing value imputation setting were aggregated using a self-weighted scheme that upweights the target modality being reconstructed.

Formally, for imputing values in a target modality *t*, we construct

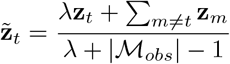

where *λ* controls the relative weight assigned to the partially observed target modality (set to *λ* = 10 in our experiments). This aggregated representation is then sent through the target modality reverse projection and decoder to reconstruct the missing values. Imputation accuracy is evaluated only at masked feature–sample entries.

We first evaluated missing value imputation under a missing completely at random (MCAR) setting, in which a fraction (*p* = 0.3) of feature values within each modality was randomly masked at test time. Figure 3a shows scatter plots comparing observed and imputed values for masked entries across modalities. Each point corresponds to a masked feature–sample pair in the test set. Imputed values closely track the observed values across a wide dynamic range, and MIMIR achieved strong correlations across all modalities, indicating accurate recovery of randomly missing values within observed modalities. Interestingly, although CNV was least accurately predicted in the missing modality scenario, it is the most accurately predicted in the missing value scenario.

**Figure 3:**
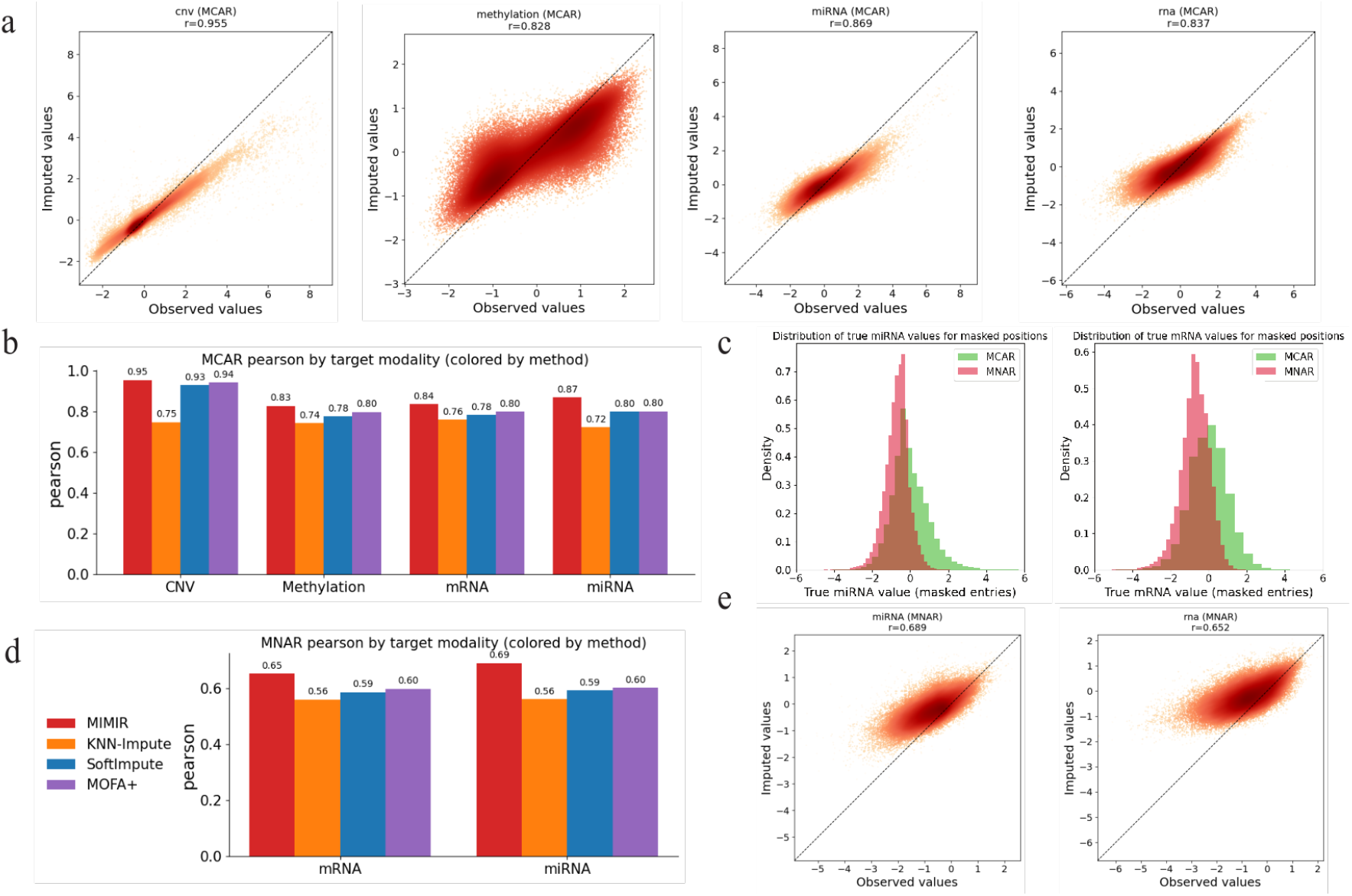
Missing value imputation under MCAR and MNAR settings. **(a)** Scatter plots comparing observed and imputed values for masked entries under MCAR masking, with each point corresponding to a masked feature–sample pair. **(b)** Comparison of MCAR imputation accuracy (Pearson correlation) with baseline methods for each target modality. **(c)** Distributions of masked values for mRNA and miRNA under MCAR and MNAR masking, showing that MNAR preferentially masks low-valued entries. **(d)** Comparison of MNAR imputation accuracy alongside baseline methods. **(e)** Scatter plots comparing observed and imputed values for masked entries under MNAR masking for mRNA and miRNA.

We then compared MIMIR’s value imputation performance against baseline methods. As baselines, we considered MOFA+, which imputes missing values by projecting corrupted samples into latent factors and reconstructing the data via learned factor loadings, a multi-omic KNN imputer, which imputes missing values based on sample similarity in the concatenated feature space, and SoftImpute, a low-rank matrix completion method applied to the concatenated multi-omic data matrix. Figure 3b summarizes Pearson correlation at masked entries for each modality across methods. Across all modalities for the MCAR setting, MIMIR consistently achieved the highest correlation, followed by MOFA+, SoftImpute and KNN-Impute. A sample of best and worse performing features in the MCAR setting are shown in Supplementary Figures 2-5.

To assess robustness to increasing levels of missingness, we varied the masking probability p across a range of values and evaluated imputation performance under MCAR (Supplementary Figure 7a). Imputation accuracy remains largely stable up to approximately 80% masking, with more pronounced degradation observed at higher levels of missingness.

In many real-world settings, missingness is structured rather than completely random and can depend on the underlying value being measured. Therefore, we next evaluated missing value imputation under a missing not at random (MNAR) setting, in which missing values preferentially affect low-valued entries within a modality. This was done to mimic abundance-dependent measurement reliability in sequencing-based assays, where low-abundance transcripts exhibit increased sampling noise and reduced detectability, approaching the effective limit of detection as demonstrated using spike-in controls. [18]. MNAR masking was applied only to mRNA and miRNA, because these sequencing-based modalities are most plausibly subject to abundance-dependent measurement effects in practice. To generate MNAR masks, we applied a ranking transform to the observed values within each modality and sampled a fraction of entries (*p* = 0.3) without replacement, with masking probabilities inversely proportional to the within-modality rank of each value.

Figure 3c compares the value distributions of masked entries under MCAR and MNAR for mRNA and miRNA, confirming that MNAR masking is strongly skewed toward lower-valued measurements. Figure 3e shows scatter plots comparing observed and imputed values for MNAR-masked entries. As expected, imputation accuracy is reduced relative to the MCAR setting due to the structured, value-dependent nature of the missingness. Nevertheless, imputed values remain well aligned with ground truth. Figure 3d summarizes Pearson correlation under MNAR across methods. While all methods experience a performance drop under MNAR, MIMIR consistently achieves higher correlation than baseline approaches for both mRNA and miRNA, indicating robustness to value-dependent missingness. Notably, the performance gap between MIMIR and baseline methods is more pronounced under MNAR than under MCAR, suggesting that MIMIR is better able to handle structured, value-dependent missingness.

As in the MCAR setting, we additionally evaluated performance across varying masking probabilities under MNAR (Supplementary Figure 7b). Compared to MCAR, performance degrades more rapidly at lower masking probabilities (p = 0.1–0.5), reflecting the increased difficulty of imputing systematically missing low-valued features. At higher masking rates, however, performance partially recovers. This likely occurs because the most difficult-to-predict features are preferentially masked at lower probabilities, and as masking increases, more predictable features are also removed.

To better understand which components of the training objective contribute to performance in both the missing modality and the missing values scenarios, we examined variants of the model with different combinations of loss terms (Supplementary Figure 1). We observe that models trained with only reconstruction or only cross-modal imputation loss perform well on their respective objectives but generalize poorly to the complementary task. Adding a contrastive alignment loss in these settings partially recovers performance on the missing objective, suggesting that the contrastive loss encourages alignment between modality-specific representations. However, when both reconstruction and cross-modal imputation losses are jointly optimized, the addition of a contrastive loss does not provide additional benefit. This indicates that the combination of reconstruction and imputation objectives already implicitly enforces alignment in the shared latent space.

### Understanding the shared space

To better understand how MIMIR integrates information across modalities, we analyzed the structure of the learned shared latent space. Because all modalities are projected into a common representation and aggregated at the sample level, the shared space provides a natural substrate for assessing how the different molecular assays contribute to, and align within, the model’s internal representation.

To characterize how individual modalities contribute to the shared latent space, we first analyzed the variance of modality-specific shared embeddings, denoted as *z*_*m*_ in the Methods section (i.e., the modality-specific embedding obtained after projection into the shared latent space), along each latent dimension (Figure 4a). We grouped dimensions according to the modality contributing the largest variance and ordered dimensions within each group by that dominant variance. This analysis indicates that the shared representation concentrates representational capacity on CNV and mRNA, as these modalities account for most of the between-sample variability in the shared space, with fewer dimensions primarily influenced by DNA methylation and none by miRNA. Although all modalities contribute to every dimension, between-sample variability in the shared space is driven primarily by CNV and mRNA, consistent with the heterogeneity of copy-number alterations and the dynamic range expected of transcriptional measurements.

**Figure 4:**
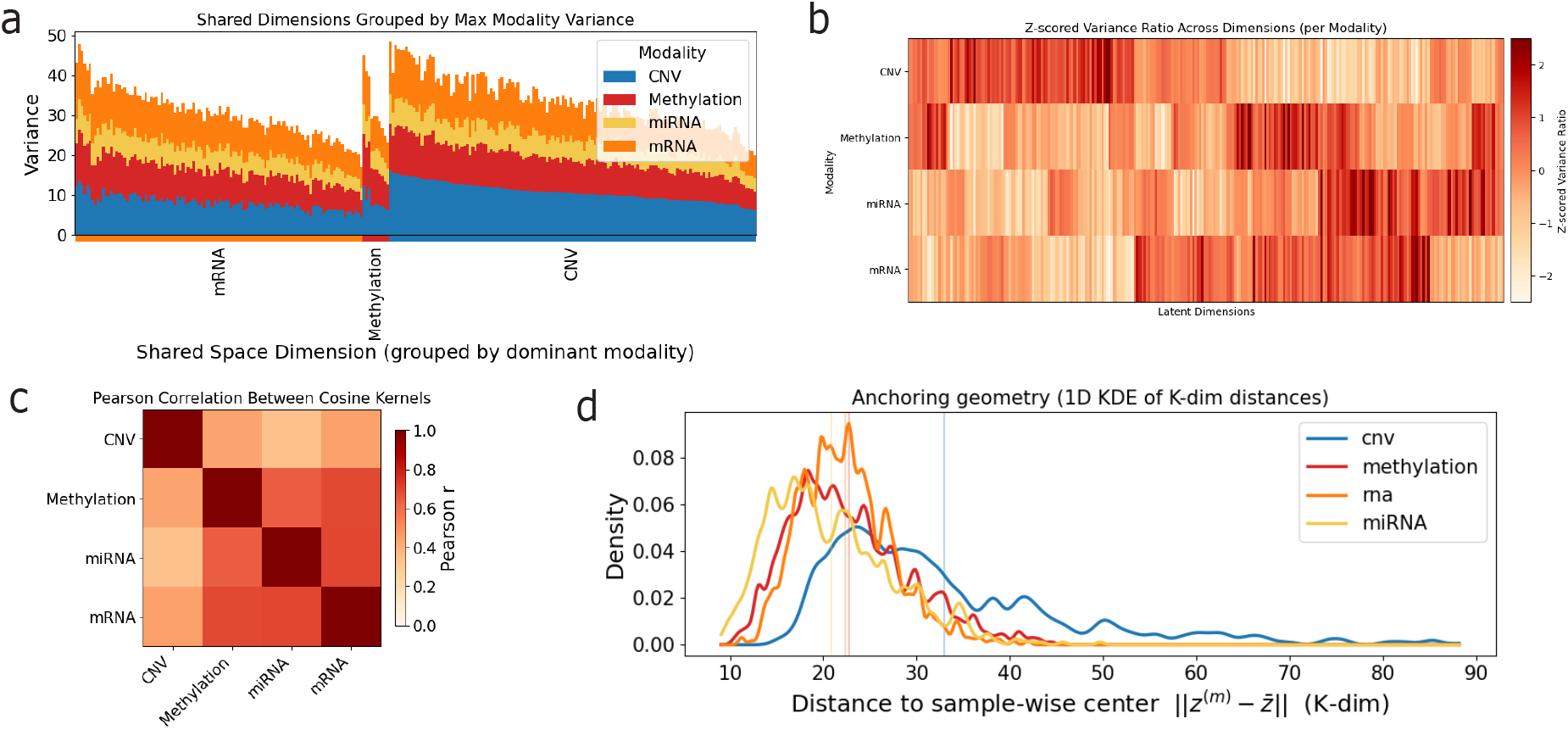
Structure of the shared latent space across modalities. **(a)** Stacked bar plot showing the variance of each modality in each shared latent dimension, after ordering dimensions by the dominant modality. **(b)** Heatmap of Z-scored relative variance contribution of each modality across shared latent dimensions. **(c)** Pairwise Pearson correlation between modality-specific similarity matrices derived from the shared embeddings, quantifying global alignment between modalities in the shared space. **(d)** Distribution of distances from each modality-specific embedding to the sample-wise centroid in the shared space. For each sample, embeddings from all observed modalities were centered by their mean, and the distance to this centroid was computed. Vertical lines indicate modality-specific mean distances.

To focus on relative patterns of variance within each modality rather than differences in absolute scale, we next analyzed standardized variance profiles across shared dimensions (Figure 4b). This transformation emphasizes which dimensions are relatively enriched or depleted for a given modality, independent of that modality’s overall variance level. Viewed in this standardized space, clear cross-modal structure emerges. The standardized variance profiles of mRNA show substantial overlap with those of miRNA and DNA methylation, indicating that these modalities tend to emphasize a shared subset of latent dimensions. In contrast, copy number variation (CNV) exhibits a largely distinct pattern, with limited overlap that is primarily restricted to a small number of dimensions shared with mRNA and methylation. This organization suggests that the shared latent space contains a core subspace jointly structured by transcriptional and epigenetic signals, alongside more separate dimensions that preferentially capture CNV-associated variation.

This observation prompted an assessment of the global alignment between modalities by comparing the similarity structure they induce over samples in the shared space (Figure 4c). For each modality, we constructed a sample–sample similarity matrix based on cosine similarity of its modality-specific shared embeddings and quantified alignment between modalities by computing pairwise Pearson correlations between these matrices. This analysis captures whether different modalities impose similar neighborhood structure over samples when represented in the shared latent space. Consistent with the standardized variance patterns in Figure 4c, strong alignment is observed between mRNA, miRNA, and DNA methylation, indicating that these modalities organize samples in a broadly concordant manner within the shared representation. In contrast, CNV exhibits substantially weaker alignment with the other modalities, reflecting a more distinct sample similarity structure that overlaps only partially with transcriptional and epigenetic signals.

Finally, we examined alignment between modality-specific embeddings at the level of individual samples by measuring each sample’s proximity to its corresponding sample-wise centroid in the shared space (Figure 4d). For each sample, modality-specific shared embeddings were centered by their mean, and the distance of each modality’s embedding to this centroid was computed. Smaller distances indicate closer agreement between a modality-specific embedding and the consensus representation inferred from all observed modalities. Consistent with the variance and alignment analyses, mRNA, miRNA, and DNA methylation embeddings tend to lie closer to the centroid, indicating stronger sample-level agreement among these modalities. In contrast, CNV embeddings are systematically farther from the centroid, reflecting greater divergence from the consensus shared representation. Together, Figures 4a–d indicate that the shared latent space is organized around a transcriptional–epigenetic core, with CNV contributing complementary structure.

## Discussion

In this work, we introduced MIMIR, a unified framework for imputing both missing modalities and missing values in multi-omic datasets through shared representation learning. By combining modality-specific autoencoders with a learned shared latent space, MIMIR accommodates arbitrary patterns of missingness while preserving cross-modal relationships that are difficult to capture with pairwise translation models or feature-level imputers. Across multiple imputation scenarios, MIMIR consistently outperformed baseline methods, demonstrating that learning a shared representation provides a practical and effective foundation for multi-omic data completion. In addition, the model remains computationally efficient, with training completing in under 30 minutes on a single GPU and scaling linearly with the number of modalities, making it practical for routine use on large multi-omic datasets.

Beyond overall performance, MIMIR exhibits systematic differences in reconstruction accuracy depending on which modalities are available. To understand these differences, we examined how imputation performance across modality subsets relates to the structure of the learned shared latent space. This analysis reveals that mRNA expression consistently provides the strongest predictive signal for reconstructing other assays, and that reconstruction of mRNA itself benefits most from the joint availability of DNA methylation and miRNA, whereas copy number variation contributes more limited gains across settings. These performance patterns are reflected in the structure of the learned shared latent space, where mRNA, miRNA, and DNA methylation exhibit overlapping regions of the shared space and therefore provide strong mutual predictive power. Meanwhile, CNV contributes more distinct and weakly aligned structure, limiting its positive impact on cross-modal reconstruction. Viewed through this lens, the prominent contribution of mRNA to imputation performance corresponds to its strong alignment with both miRNA and DNA methylation in the shared latent space. This is consistent with the role of gene expression as an integrative readout of epigenetic and post-transcriptional regulation. In contrast, CNV contributes variance along more weakly aligned dimensions, reflecting the fact that copy number alterations are further upstream and do not uniformly propagate to downstream molecular layers. Importantly, these patterns emerge naturally from the model without imposing explicit assumptions about modality hierarchy, indicating that MIMIR captures intrinsic biological dependencies across molecular layers rather than treating modalities as interchangeable inputs.

At a high level, MIMIR shares conceptual similarities with latent factor models such as MOFA+, in that both approaches seek to represent multiple molecular modalities within a common low-dimensional space. However, MIMIR differs in several important respects. While MOFA+ relies on a linear generative model with shared latent factors learned jointly across modalities, MIMIR employs nonlinear modality-specific encoders and decoders, enabling it to capture more complex, non-additive relationships within and across molecular layers. MIMIR is also related to recent deep learning frameworks for bulk multi-omic integration, such as Integrate Any Omics and DeepIMV, which learn joint representations across modalities but are primarily designed for downstream prediction tasks rather than imputation [19, 20]. In addition, MIMIR is conceptually related to single-cell integration methods, which often rely on modality-specific generative assumptions or pairwise translation between modalities, and in some cases combine both approaches [15, 14, 21]. However, across both bulk and single-cell settings, existing approaches are not designed to jointly address multiple forms of missingness. In contrast, MIMIR adopts a modality-agnostic representation learning framework and is specifically designed for bulk multi-omic imputation, addressing two distinct but common challenges in multi-omic studies: missing entire modalities and missing values within observed modalities. By unifying these imputation tasks within a single shared-representation framework, MIMIR supports flexible inference from arbitrary subsets of available data without requiring separate models or procedures for modality-level and feature-level completion.

In parallel, a separate line of work has focused on imputing missing epigenomic measurements. Methods such as ChromImpute, PREDICTD, and Avocado leverage shared structure across sample types, assays, and genomic coordinates, often framing the problem as tensor completion over a fixed genomic axis [22, 23, 24]. Because these approaches are inherently tied to genomic position and rely on a shared coordinate system, they do not generalize to data modalities that lack such a structure and therefore address a slightly different class of imputation problems than sample-level multi-omic imputation. MIMIR instead operates in this broader setting, enabling imputation across more heterogeneous modalities and missingness patterns within a unified framework.

Several directions for future work follow naturally from this framework. Extending MIMIR to additional molecular modalities and to datasets with higher degrees of incompleteness will further test its scalability and robustness. In particular, integrating additional modalities such as proteomics or imaging-based assays, which often exhibit structured or detection-limited missingness, may further highlight the benefits of explicitly modeling non-random missingness in multi-omic data. In addition, exploring alternative aggregation strategies, such as learned or context-dependent weighting of modalities, may improve imputation accuracy in settings where data quality or informativeness varies across assays. A further limitation of the current framework is that modalities are treated as unordered feature vectors, without explicitly incorporating genomic coordinate structure. While this design enables MIMIR to operate across heterogeneous data types, incorporating genomic positional information as an additional inductive bias may improve imputation accuracy for modalities that have a genomic axis. Finally, leveraging larger models pretrained on single-modality datasets before joint fine-tuning may improve representations when multi-omic sample sizes are limited.

The MIMIR framework is available as open-source software, and users can readily retrain models from scratch on their own multi-omic datasets, enabling adaptation to different feature spaces, modality combinations, and study designs.

## Materials and methods

### Data

We used publicly available pan-cancer multi-omic data derived from The Cancer Genome Atlas (TCGA, https://www.cancer.gov/ccg/research/genome-sequencing/tcga). To define a consistent sample and feature schema across molecular modalities, we relied on the MLOmics pan-cancer dataset, which provides harmonized sample identifiers and feature definitions across multiple omics layers [25]. Importantly, we used MLOmics only to determine shared sample identifiers and feature definitions. All molecular measurements were reloaded and reprocessed directly from the original TCGA Xena data files to avoid inheriting any imputed values or downstream feature selection applied in the MLOmics pipeline [26].

We considered four molecular modalities: mRNA expression, microRNA (miRNA) expression, DNA methylation, and copy number variation (CNV). mRNA expression values were obtained as log_2_(*TPM* + 0.001)–transformed RSEM estimates from TCGA RNA-seq data. miRNA expression values were obtained from TCGA miRNA sequencing data and log-transformed in the same manner. DNA methylation data consisted of beta values, defined as the proportion of methylated signal at a given CpG locus, and was generated using the Illumina HumanMethylation450 array. CNV profiles were represented as gene-level copy number estimates derived from GISTIC2 [27] calls. For each modality, features were restricted to those present in the MLOmics pan-cancer dataset to ensure consistent dimensionality across samples, but no additional filtering, ranking, or imputation was applied. The number of features for each modality is shown in Figure 1b. Approximately 8,000 samples had all four modalities available and were used for model training. All features within each modality were Z-score normalized across samples. Samples were split at the individual level into training, validation, and test sets, with all modalities corresponding to a given individual assigned to the same split.

Although the MLOmics pan-cancer dataset contains complete multi-omic profiles at the modality level by construction, some feature-level sparsity and missingness remain within individual modalities. During model training and evaluation, both modality-level and feature-level missingness were introduced synthetically, enabling controlled assessment of cross-modal reconstruction and imputation performance.

### Modeling phase 1: modality autoencoders

The first phase of MIMIR consists of training an independent autoencoder for each molecular modality. This stage serves two purposes: (i) to learn compact, modality-specific latent representations that capture the dominant structure within each data type, and (ii) to initialize encoders and decoders that are robust to feature-level missingness, which later facilitates cross-modal integration and imputation. Importantly, during this phase, no information is shared across modalities. Each autoencoder is trained entirely independently.

#### Model architecture

For each modality, *m* ∈ {mRNA, miRNA, methylation

, CNV }, we train a feedforward autoencoder consisting of an encoder *f*_*m*_(·) and a decoder *g*_*m*_(·). The encoder maps an input feature vector 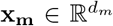 to a lower-dimensional latent representation 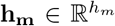, and the decoder reconstructs the input from this latent space:

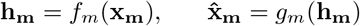

Both encoder and decoder are implemented as multilayer perceptrons with ReLU activations. The depth and width of each autoencoder are modality-specific, reflecting differences in dimensionality and signal structure across data types. In practice, we used a single hidden layer for all modalities, with latent dimensionalities of 512 for mRNA, 256 for methylation and CNV, and 128 for miRNA (Figure 1b). Dropout was applied after hidden activations to regularize training.

To enable downstream feature-level imputation, the autoencoders are trained using explicit feature masking. With probability *p* = 0.15, individual feature values are replaced with a sentinel value (zero in all experiments). The model is then trained to reconstruct the original, unmasked input. This approach has been shown to learn robust representations in the presence of missing or corrupted inputs [28]. Moreover, it allows the autoencoder to explicitly learn to infer missing values from observed features within the same modality. The masking operation is applied only during training and is disabled at test time.

#### Training procedure

Each modality-specific autoencoder is trained using mean squared error (MSE) reconstruction loss, with careful handling of both truly missing values and artificially masked entries. Truly missing values (NaNs arising from assay limitations or biological sparsity) are excluded from all loss calculations and are replaced with the modality’s sentinel value before being passed through the network. Artificially masked values introduced during denoising are tracked explicitly and used to compute a masked reconstruction loss.

Specifically, for each batch, we compute (i) an overall reconstruction loss over all observed (non-missing) features, and (ii) a masked reconstruction loss restricted to artificially masked positions. These two terms are combined as a weighted sum:

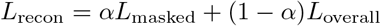

where *α* = 0.5 in all experiments, placing equal emphasis on accurate global reconstruction and masked-value imputation.

At the end of Phase 1, the encoder and decoder weights for each modality are saved and used to initialize the shared-representation model in Phase 2.

### Modeling phase 2: shared representations

In Phase 2, we integrate the pretrained modality-specific autoencoders into a unified model that learns a shared latent representation across molecular layers (Figure 1c). The goal of this phase is twofold: (i) to align representations from different modalities into a common latent space that captures cross-modal dependencies, and (ii) to enable imputation of both missing modalities and missing feature values within observed modalities.

#### Model architecture

Using the modality-specific encoders *f*_*m*_(.) trained in Phase 1, each observed modality is mapped to its latent representation *h*_*m*_. To enable cross-modal interaction, each *h*_*m*_ is projected into a shared latent space of fixed dimensionality via a learnable, modality-specific projection head:

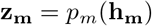

All projection heads map into the same shared space of dimension 256, allowing representations from different modalities to be directly compared and combined. Learning a shared latent representation across modalities is a common strategy in multi-modal representation learning [29, 30]. When multiple modalities are observed for a sample, the shared representation is formed by averaging the corresponding projected embeddings:

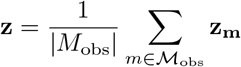

where *M*_obs_ denotes the set of observed modalities for that sample. This aggregation produces a single shared latent representation regardless of which subset of modalities is available. To reconstruct modality-specific data from the shared space, inverse projection heads *r*_*m*_(·) map the aggregated shared representation **z** back to modality-specific latent spaces, after which the pretrained decoders *g*_*m*_(·) reconstruct the original features, 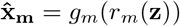. This structure allows a modality to be reconstructed either from its own shared embedding or from a shared representation inferred from other observed modalities.

#### Training procedure

The shared-representation model is trained end-to-end by fine-tuning the pretrained modality-specific encoders and decoders together with the projection and inverse-projection heads introduced in Phase 2.

During training, both feature-level and modality-level missingness are explicitly simulated. Feature-level missingness is introduced by randomly masking a fraction of observed feature values within each modality and replacing them with a sentinel value, following the masked reconstruction scheme described in Phase 1. Values that are truly missing from the data are excluded from masking and from all loss computations. Training is performed using samples with all modalities observed. However, to simulate missing modalities, entire modalities are randomly dropped with a probability of 0.4 on a per-sample basis during training. When a modality is dropped, it is omitted from the forward pass, and the shared representation is computed by averaging the projected embeddings of the remaining observed modalities. This procedure exposes the model to arbitrary subsets of available modalities and encourages robustness to modality-level missingness at test time. In addition to this stochastic modality dropout, cross-modal imputation is trained using a leave-one-modality-out objective applied to all observed modalities, as described below.

Training jointly optimizes three objectives: masked reconstruction within modalities, contrastive alignment across modalities, and cross-modal imputation.

##### Masked Reconstruction Loss

For each observed modality, we apply the same masked reconstruction objective used in Phase 1. A subset of observed feature values is randomly masked and replaced with a sentinel value, and reconstruction error is computed using mean squared error over valid (non-missing) entries. The loss combines reconstruction over all observed features with reconstruction restricted to masked features, encouraging robustness to feature-level missingness while preserving overall reconstruction fidelity. Values that are truly missing from the data are excluded from all loss computations.

##### Contrastive Alignment Loss

To align representations across modalities, we apply a contrastive loss to the shared embeddings. For a given sample, shared embeddings derived from different modalities of the same sample are treated as positive pairs, while embeddings from different samples in the same batch act as negatives. Similarity is computed using cosine similarity with a temperature parameter *τ* = 0.1, which controls the sharpness of the similarity distribution in the contrastive objective, and the loss is implemented using a cross-entropy objective over the batch. Formally, let ℳ_*obs*_ be the set of observed modalities in a batch and let 𝒫 be the set of ordered modality pairs (*m, m*^′^) with *m* ≠ *m*^′^. For each ordered pair (*m, m*^′^) define a cosine similarity matrix, 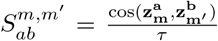 for *a, b* ∈ {1, …, *B*} where *B* is the batch size. The contrastive loss is then

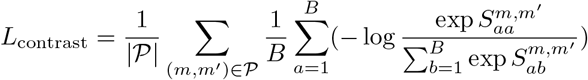

This objective follows the InfoNCE formulation used in contrastive representation learning [31]

##### Cross-Modal Imputation Loss

To explicitly train the model to perform cross-modal imputation, we adopt a leave-one-modality-out reconstruction objective. For each sample and for each modality *m* that is present, the shared representation is computed by averaging the shared embeddings of all other available modalities. This aggregated representation is then mapped back through the inverse projection and decoder for modality *m* to reconstruct its features. Reconstruction error for the target modality is penalized using mean squared error over valid features. This loss encourages the shared latent space to retain sufficient information to support modality-level imputation under arbitrary missingness patterns.

##### Overall Objective

The full training objective is a weighted sum of the three loss components:

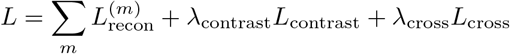

Here, the reconstruction loss is summed over all observed modalities for each sample, while the cross-modal imputation loss *L*_cross_ is computed as the average reconstruction error across all target modalities, where each target modality is reconstructed from the remaining available modalities in a leave-one-modality-out fashion. Unless otherwise specified, we set *λ*_contrast_ = 1 and *λ*_cross_ = 1 in all experiments.

Optimization is performed using the Adam optimizer with an initial learning rate of 3 × 10^−4^ and weight decay of 10^−5^. Due to the compact model architecture, the combined training time for both phases was under 30 minutes on a single NVIDIA GeForce RTX 2080 GPU. The model scales linearly with the number of modalities, as each modality is encoded independently before integration into the shared latent space. Training proceeds in mini-batches, with all available modalities for a given individual processed together. Model selection is based on validation loss, and all reported results are obtained from held-out test samples.

### Baseline Methods

#### Missing modality imputation baselines

MOFA+ is a probabilistic latent factor model that jointly embeds multiple omic modalities into a shared low-dimensional representation via Bayesian matrix factorization [16]. We trained a single global MOFA+ model on the training samples using all modalities. Because MOFA+ does not provide a built-in procedure for missing-modality prediction, we performed post hoc inference by projecting scenario samples into the learned latent space using observed modalities and reconstructing the missing modality from the inferred factors using the corresponding loading matrix. When multiple modalities were observed, factor projections were obtained using a least-squares multi-view projection where information across all available views are aggregated. This procedure essentially uses the learned latent representation to generate predictions for the missing modality.

TOBMI* is a k-nearest-neighbor–based trans-omics imputation method that reconstructs a missing modality by borrowing information from similar samples identified in the observed modalities [17]. For each missing-modality scenario, we identified nearest neighbors among training samples using cosine distance in the concatenated feature space of the observed modalities and reconstructed the missing modality via distance-weighted regression. The number of neighbors was set to the square root of the number of training samples. In our implementation, we replaced the Mahalanobis distance used in the original TOBMI formulation with cosine distance due to the absence of an official implementation and the computational challenges associated with estimating and inverting high-dimensional covariance matrices in this setting.

#### Missing values imputation baselines

The KNN-Imputer concatenates features across selected modalities and imputes missing entries based on nearest neighbors in the resulting feature space, using the standard nan-aware Euclidean distance. The imputer was fit on training samples and applied to corrupted validation and test data with the same feature layout.

SoftImpute is a low-rank matrix completion method that performs iterative singular value thresholding on a concatenated multi-omic data matrix [8]. We applied SoftImpute directly to corrupted data matrices containing missing values. Hyperparameters controlling the maximum rank and singular value shrinkage were selected on the validation set MCAR and then fixed for evaluation on the test set.

MOFA+ was also used as a baseline for missing value imputation. Using the global model trained on all modalities, corrupted samples were projected into the latent space using observed entries (via the same least-squares multi-view projection as in the missing-modality setting) and reconstructed via the learned loading matrices. Imputed values were taken only at missing entries while preserving observed values.

## Availability and Implementation

The code and data used in this paper is available at: https://github.com/Noble-Lab/MIMIR

## Acknowledgements

This work is supported in part by funds from the National Institutes of Health (U01 HG013198).

## Author contributions

Conceptualization: A.N., C.M., W.S.N. Data curation: A.N., C.M. Funding acquisition, Supervision, Resources: W.S.N. Investigation: A.N., C.M., W.S.N. Methodology: A.N., W.S.N. Software, Visualization: A.N. Writing: A.N., W.S.N.

## Interests statement

No competing interests.

## Notes

### Competing Interest Statement

The authors have declared no competing interest.

### Summary of Updates

The MOFA baseline is now used for both missing modality and missing value imputation. We also now include a Supplementary Information document to complement the main text.

## References

[1] Arno van Hilten, Steven A Kushner, Manfred Kayser, M Arfan Ikram, Hieab HH Adams, Caroline CW Klaver, Wiro J Niessen, and Gennady V Roshchupkin. Gennet framework: interpretable deep learning for predicting phenotypes from genetic data. Communications Biology, 4(1):1094, 2021.

[2] Stephen B Montgomery, Jonathan A Bernstein, and Matthew T Wheeler. Toward transcriptomics as a primary tool for rare disease investigation. Molecular Case Studies, 8(2):a006198, 2022.

[3] Craig Smail and Stephen B Montgomery. Rna sequencing in disease diagnosis. Annual Review of Genomics and Human Genetics, 25, 2024.

[4] Anahita Samih, Maurício Alexander de Moura Ferreira, and Zoran Nikoloski. Gene expression and protein abundance: Just how associated are these molecular traits? Biotechnology Advances, page 108720, 2025.

[5] Ruochen Liu, Erhu Zhao, Huijuan Yu, Chaoyu Yuan, Muhammad Nadeem Abbas, and Hongjuan Cui. Methylation across the central dogma in health and diseases: new therapeutic strategies. Signal Transduction and Targeted Therapy, 8(1):310, 2023.

[6] Žiga Avsec, Vikram Agarwal, Daniel Visentin, Joseph R Ledsam, Agnieszka Grabska-Barwinska, Kyle R Taylor, Yannis Assael, John Jumper, Pushmeet Kohli, and David R Kelley. Effective gene expression prediction from sequence by integrating long-range interactions. Nature methods, 18(10):1196–1203, 2021.

[7] Žiga Avsec, Natasha Latysheva, Jun Cheng, Guido Novati, Kyle R Taylor, Tom Ward, Clare Bycroft, Lauren Nicolaisen, Eirini Arvaniti, Joshua Pan, et al. Advancing regulatory variant effect prediction with alphagenome. Nature, 649(8099):1206–1218, 2026.

[8] Rahul Mazumder, Trevor Hastie, and Robert Tibshirani. Spectral regularization algorithms for learning large incomplete matrices. The Journal of Machine Learning Research, 11:2287–2322, 2010.

[9] Ricard Argelaguet, Britta Velten, Damien Arnol, Sascha Dietrich, Thorsten Zenz, John C Marioni, Florian Buettner, Wolfgang Huber, and Oliver Stegle. Multiomics factor analysis—a framework for unsupervised integration of multi-omics data sets. Molecular Systems Biology, 14(6):e8124, 2018.

[10] Daniel J Stekhoven and Peter Bühlmann. Missforest—non-parametric missing value imputation for mixed-type data. Bioinformatics, 28(1):112–118, 2012.

[11] Md Istiaq Ansari, Khandakar Tanvir Ahmed, and Wei Zhang. Optimizing multi-omics data imputation with nmf and gan synergy. Bioinformatics, 40(11):btae674, 2024.

[12] Xiang Zhou, Hua Chai, Huiying Zhao, Ching-Hsing Luo, and Yuedong Yang. Imputing missing rna-sequencing data from dna methylation by using a transfer learning–based neural network. GigaScience, 9(7):giaa076, 2020.

[13] Soyeon Kim, Hyun Jung Park, Xiangqin Cui, and Degui Zhi. Collective effects of long-range dna methylations predict gene expressions and estimate phenotypes in cancer. Scientific Reports, 10(1):3920, 2020.

[14] Kevin E Wu, Kathryn E Yost, Howard Y Chang, and James Zou. Babel enables cross-modality translation between multiomic profiles at single-cell resolution. Proceedings of the National Academy of Sciences, 118(15):e2023070118, 2021.

[15] Ran Zhang, Laetitia Meng-Papaxanthos, Jean-Philippe Vert, and William Stafford Noble. Semi-supervised single-cell cross-modality translation using polarbear. In International Conference on Research in Computational Molecular Biology, pages 20–35. Springer, 2022.

[16] Ricard Argelaguet, Damien Arnol, Danila Bredikhin, Yonatan Deloro, Britta Velten, John C Marioni, and Oliver Stegle. Mofa+: a statistical framework for comprehensive integration of multi-modal single-cell data. Genome Biology, 21(1):111, 2020.

[17] Xuesi Dong, Lijuan Lin, Ruyang Zhang, Yang Zhao, David C Christiani, Yongyue Wei, and Feng Chen. Tobmi: trans-omics block missing data imputation using a k-nearest neighbor weighted approach. Bioinformatics, 35(8):1278–1283, 2019.

[18] Sarah A Munro, Steven P Lund, P Scott Pine, Hans Binder, Djork-Arné Clevert Ana Conesa, Joaquin Dopazo, Mario Fasold, Sepp Hochreiter, Huixiao Hong, et al. Assessing technical performance in differential gene expression experiments with external spike-in rna control ratio mixtures. Nature communications, 5(1):5125, 2014.

[19] Shihao Ma, Andy GX Zeng, Benjamin Haibe-Kains, Anna Goldenberg, John E Dick, and Bo Wang. Moving towards genome-wide data integration for patient stratification with integrate any omics. Nature Machine Intelligence, 7(1):29–42, 2025.

[20] Changhee Lee and Mihaela Van der Schaar. A variational information bottleneck approach to multiomics data integration. In International Conference on Artificial Intelligence and Statistics, pages 1513–1521. PMLR, 2021.

[21] Tal Ashuach, Mariano I Gabitto, Rohan V Koodli, Giuseppe-Antonio Saldi, Michael I Jordan, and Nir Yosef. Multivi: deep generative model for the integration of multimodal data. Nature methods, 20(8):1222–1231, 2023.

[22] Jason Ernst and Manolis Kellis. Large-scale imputation of epigenomic datasets for systematic annotation of diverse human tissues. Nature biotechnology, 33(4):364–376, 2015.

[23] Jacob Schreiber, Timothy Durham, Jeffrey Bilmes, and William Stafford Noble. Avocado: a multi-scale deep tensor factorization method learns a latent representation of the human epigenome. Genome biology, 21(1):81, 2020.

[24] Timothy J Durham, Maxwell W Libbrecht, J Jeffry Howbert, Jeff Bilmes, and William Stafford Noble. Predictd parallel epigenomics data imputation with cloud-based tensor decomposition. Nature communications, 9(1):1402, 2018.

[25] Ziwei Yang, Rikuto Kotoge, Xihao Piao, Zheng Chen, Lingwei Zhu, Peng Gao, Yasuko Matsubara, Yasushi Sakurai, and Jimeng Sun. Mlomics: Cancer multi-omics database for machine learning. Scientific Data, 12(1):913, 2025.

[26] Mary J Goldman, Brian Craft, Mim Hastie, Kristupas Repečka, Fran McDade, Akhil Kamath, Ayan Banerjee, Yunhai Luo, Dave Rogers, Angela N Brooks, et al. Visualizing and interpreting cancer genomics data via the xena platform. Nature Biotechnology, 38(6):675–678, 2020.

[27] Craig H Mermel, Steven E Schumacher, Barbara Hill, Matthew L Meyerson, Rameen Beroukhim, and Gad Getz. Gistic2. 0 facilitates sensitive and confident localization of the targets of focal somatic copy-number alteration in human cancers. Genome Biology, 12(4):R41, 2011.

[28] Pascal Vincent, Hugo Larochelle, Yoshua Bengio, and Pierre-Antoine Manzagol. Extracting and composing robust features with denoising autoencoders. In Proceedings of the 25th International Conference on Machine learning, pages 1096–1103, 2008.

[29] Tadas Baltrušaitis, Chaitanya Ahuja, and Louis-Philippe Morency. Multimodal machine learning: A survey and taxonomy. IEEE Transactions on Pattern Analysis and Machine Intelligence, 41(2):423–443, 2018.

[30] Nitish Srivastava and Russ R Salakhutdinov. Multimodal learning with deep boltzmann machines. Advances in Neural Information Processing Systems, 25, 2012.

[31] Aaron van den Oord, Yazhe Li, and Oriol Vinyals. Representation learning with contrastive predictive coding. arXiv preprint arXiv:1807.03748, 2018.

