## Supplemental Information for "Unified imputation of missing data modalities and features in multi-omic data via shared representation learning"

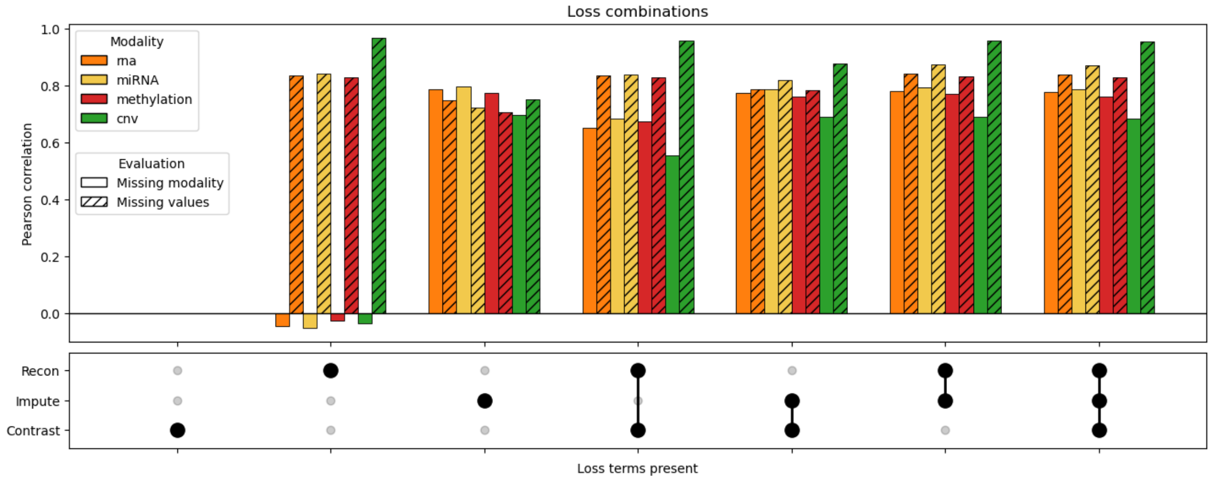


Supp. Fig. 1: **Effect of loss function combinations on imputation performance.** Comparison of Pearson correlation across modalities (mRNA, miRNA, methylation, and CNV) for different combinations of reconstruction, imputation, and contrastive loss terms. Bars show performance for each modality, with solid colors indicating missing modality imputation and hatched bars indicating missing value imputation. The bottom panel indicates which loss terms are active in each configuration.

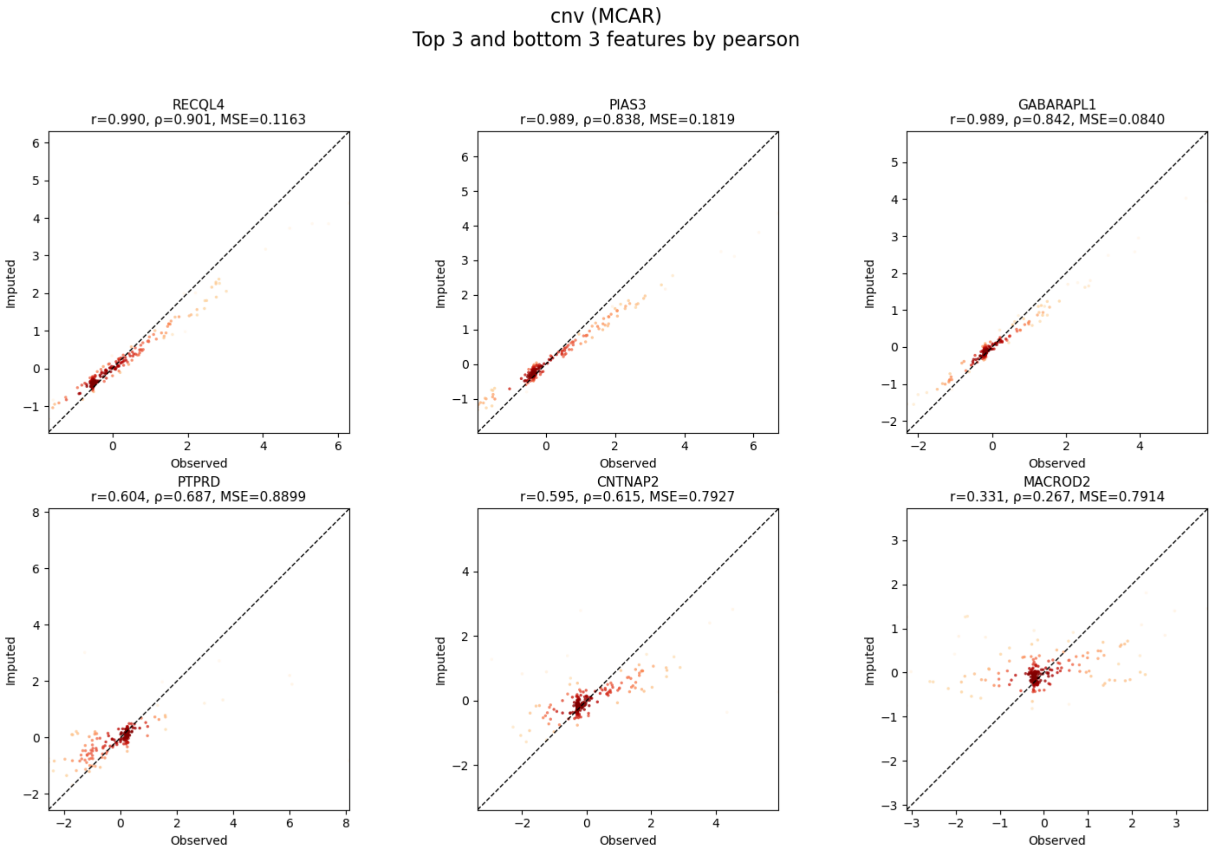


Supp. Fig. 2: **Examples of best- and worst-predicted CNV features (MCAR masking).** Scatter plots of observed versus predicted values for the top three and bottom three CNV features ranked by Pearson correlation. Each panel corresponds to a single feature, with reported correlation (r), p-value, and mean squared error (MSE). Dashed lines indicate the identity line.

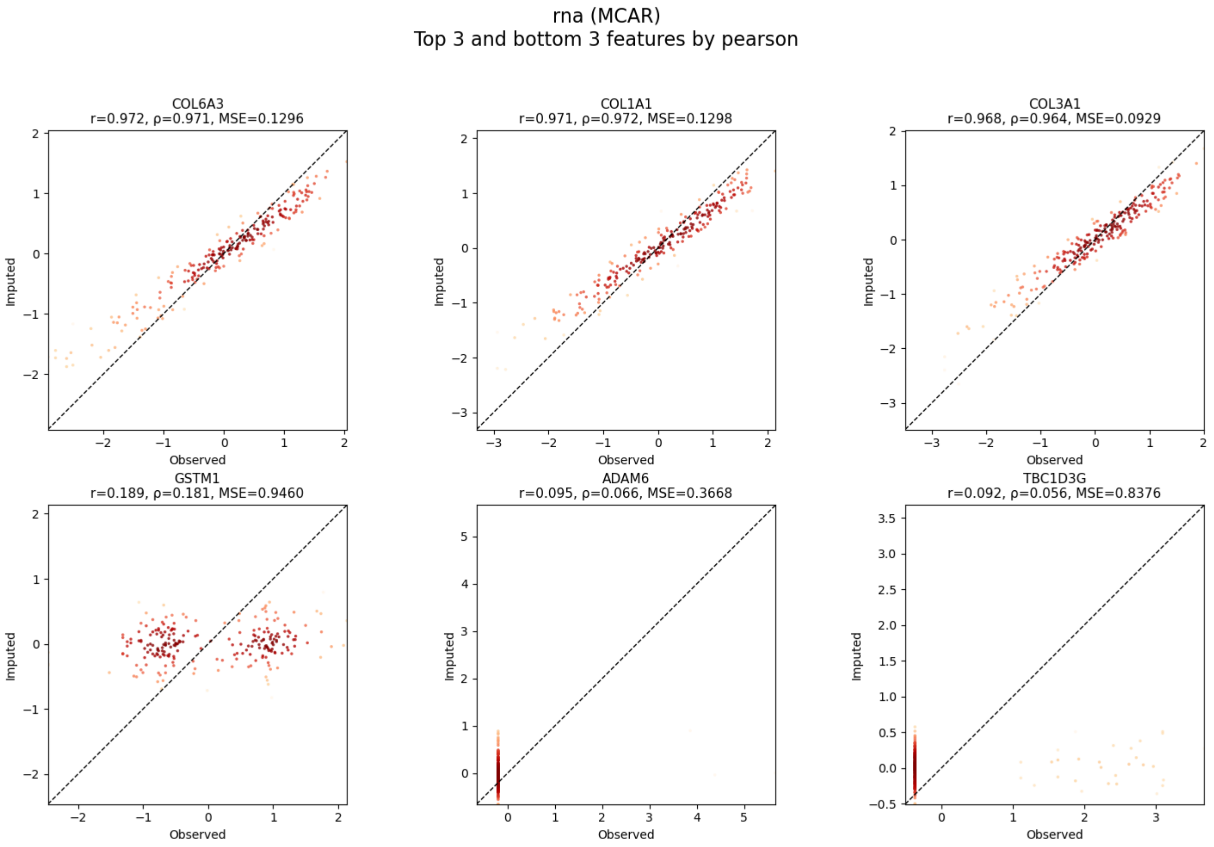


Supp. Fig. 3: **Examples of best- and worst-predicted mRNA features (MCAR masking).** Scatter plots of observed versus predicted values for the top three and bottom three mRNA features ranked by Pearson correlation. Each panel shows one gene, with correlation (r), p-value, and MSE annotated. Dashed lines indicate the identity line.

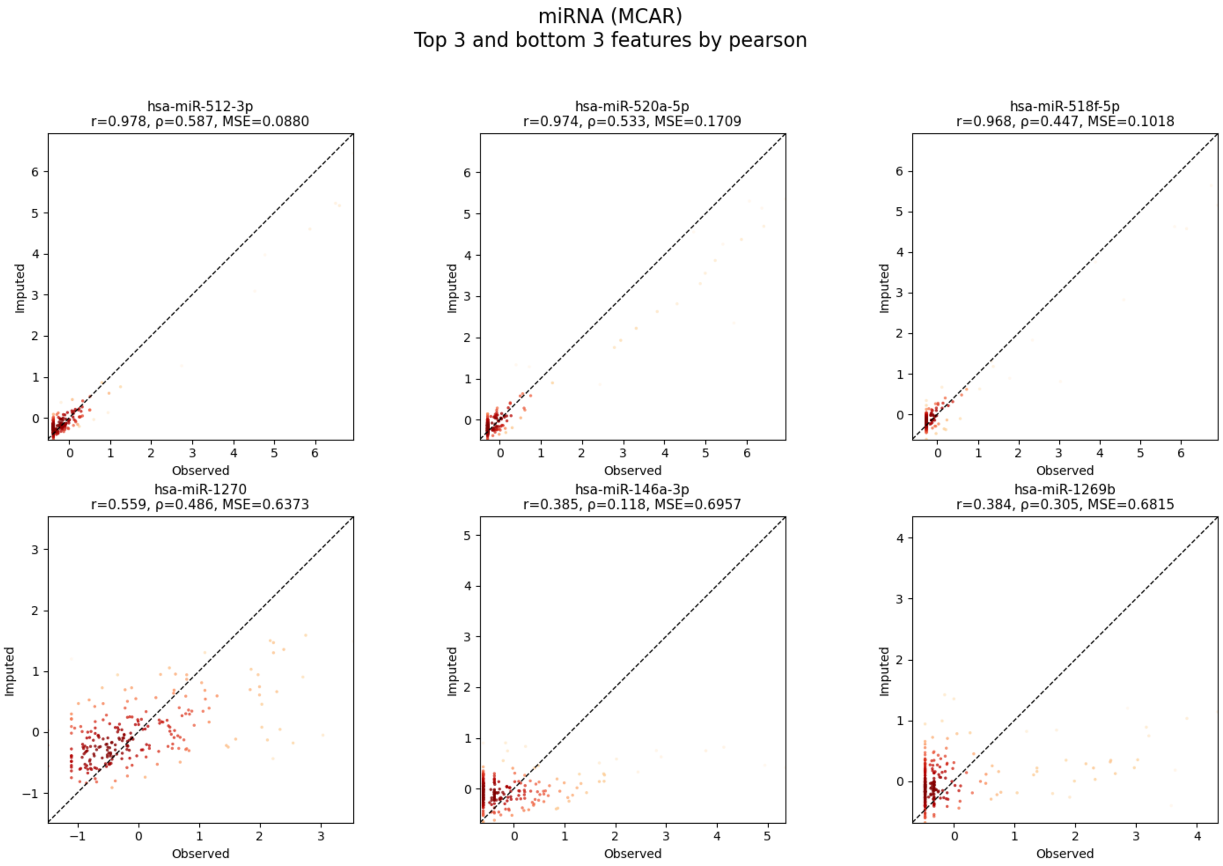


Supp. Fig. 4: **Examples of best- and worst-predicted mRNA features (MCAR masking).** Scatter plots of observed versus predicted values for the top three and bottom three mRNA features ranked by Pearson correlation. Each panel shows one gene, with correlation (r), p-value, and MSE annotated. Dashed lines indicate the identity line.

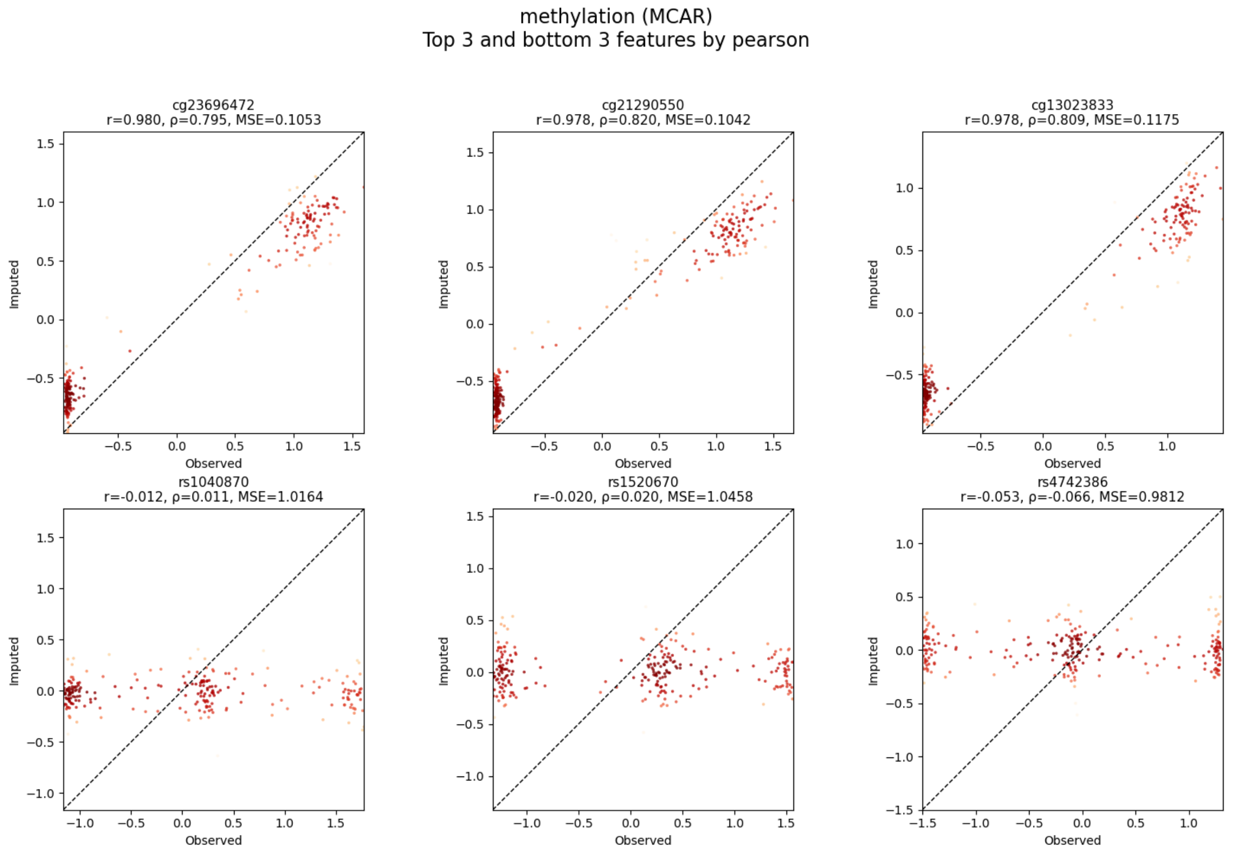


Supp. Fig. 5: **Examples of best- and worst-predicted methylation features (MCAR masking).** Scatter plots of observed versus predicted values for the top three and bottom three mRNA features ranked by Pearson correlation. Each panel shows one gene, with correlation (r), p-value, and MSE annotated. Dashed lines indicate the identity line.

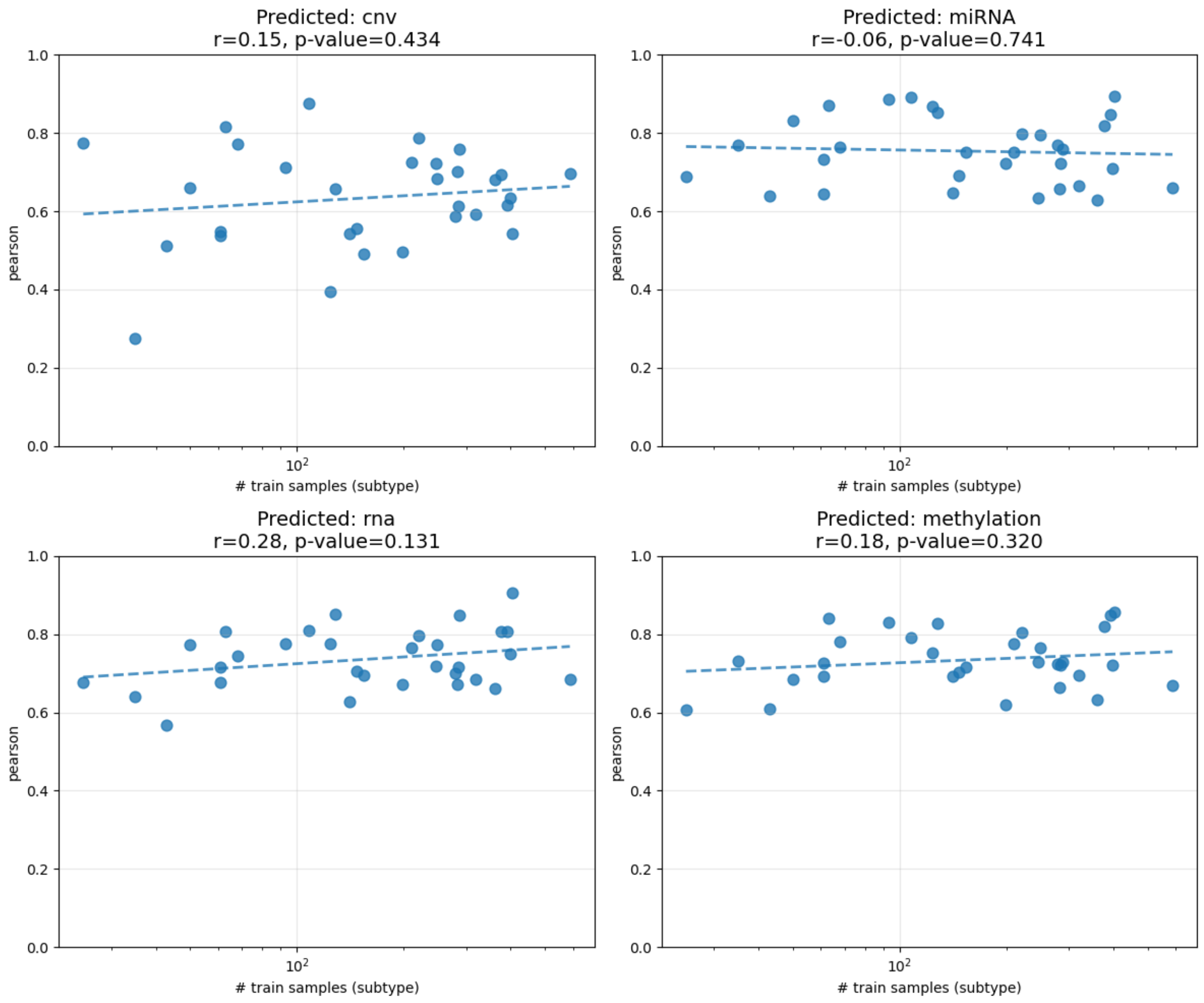


Supp. Fig. 6: **Relationship between training sample size and imputation performance across modalities.** Pearson correlation is plotted against the number of training samples for each subtype across CNV, miRNA, mRNA, and methylation. Each point represents a subtype-specific model, and dashed lines show linear trends. Reported correlation coefficients (r) and p-values quantify the relationship between sample size and performance.

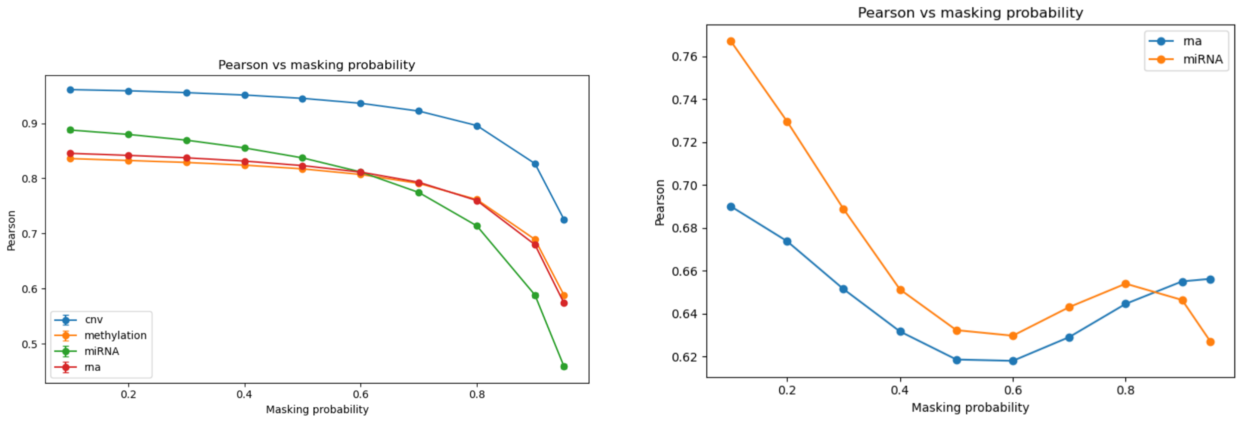


(a) MCAR

(b) MNAR

Supp. Fig. 7: **Effect of masking probability on imputation performance under MCAR and MNAR settings.** **(a)** Pearson correlation as a function of masking probability under MCAR masking. **(b)** Pearson correlation under MNAR masking, where low-valued entries are preferentially masked. Performance is shown for mRNA, miRNA, methylation, and CNV, demonstrating differing robustness of modalities to increasing missingness.
